# Improved Prediction of Bacterial Type VI Secretion Effector Proteins Using an Integrated Convolutional Neural Network Model Combining N-terminal Signal Sequences, Evolutionary Information and Pre-Trained Protein Language Features

**DOI:** 10.1101/2025.03.07.642067

**Authors:** Yueming Hu, Minqing Yan, Yanyan Zhu, Haoyu Chao, Sida Li, Qinyang Ni, Yanshi Hu, Enyan Liu, Liya Liu, Yifan Chen, Zeya Zhou, Yuhao Chen, Shilong Zhang, Yejun Wang, Cong Feng, Ming Chen

## Abstract

Type VI secretion system effectors (T6SEs) are crucial for bacterial pathogenicity, making their accurate identification essential for understanding bacterial virulence mechanisms. This study analyzed the differences in amino acid composition of N-terminal signal sequences between T6SEs and non-T6SEs, uncovering distinct positional amino acid preferences in T6SEs. Using a combination of unsupervised and supervised analysis, we evaluated feature encoding methods and developed T6CNN, an ensemble model that integrates N-terminal signal sequences, evolutionary information, and pre-trained protein language features for T6SE prediction. T6CNN demonstrated outstanding performance in independent testing, outperforming existing tools with a 7.9% accuracy increase (to 0.953), a 13.2% sensitivity improvement (to 0.964), and a 6.6% specificity enhancement (to 0.951). The T6CNN model offers a reliable and accurate solution for T6SE prediction, with significant potential to advance research on bacterial pathogenicity.

**Importance:** This study introduces T6CNN, a new computational model for identifying Type VI secretion system effectors used by harmful bacteria. By analyzing early protein sequences, T6CNN uncovers unique features that reliably distinguish effector proteins. Integrating evolutionary data and pre-trained protein language features, the model outperforms existing methods in accuracy, sensitivity, and specificity. This enhanced prediction tool deepens our understanding of bacterial infection mechanisms and offers researchers a valuable resource for pinpointing key virulence factors. Ultimately, T6CNN may help drive the development of more targeted antibacterial treatments and strategies to combat bacterial diseases.

## Introduction

Bacteria employ multiple macromolecular nanomachines to secrete proteins and other molecules. To date, over a dozen protein secretion systems have been identified in Gram-negative bacteria, including two inner membrane-spanning export pathways (Sec and Tat) and type I-IX secretion systems (T1SSs–T9SSs) (1). The type VI secretion system (T6SS) was first functionally characterized in 2006 in *Vibrio cholerae* (2). T6SSs affect eukaryotic and bacterial cells by releasing substrates known as effectors (T6SEs), which are critical for bacterial pathogenicity, competition and survival (3, 4).

The T6SS employs diverse strategies for substrate transport, with effector proteins broadly categorized as “specialized” effectors and “cargo” effectors (5, 6). “Specialized” effectors are homologous to T6SS structural components (Hcp, VgrG, and PAAR) and are covalently linked to their C-terminus (7). In contrast, “cargo” effectors interact with T6SS core components either directly or indirectly through adapter proteins (8, 9). These effectors contain specific motifs and domains, including Rhs/YD repeat sequences, N-terminal PAAR domains, C-terminal effector domains, TTR domains, MIX motifs, and FIX domains of unknown function, which are essential for identifying and characterizing T6SEs (10–13).

Predicting and identifying T6SEs, as well as elucidating their pathogenic mechanisms, are crucial for understanding bacterial interactions with hosts and other bacteria and for developing novel antibacterial drugs (14). Similar to effectors in other translocation systems, T6SEs exhibit substantial diversity, making the experimental identification of novel T6SEs a significant challenge (6). Since the introduction of the first machine learning algorithm for T6SE prediction in 2018, only a few computational models have been developed for this purpose (15–18). Bastion6, the first machine learning-based T6SE prediction tool, utilizes sequence features, evolutionary data, and physicochemical properties to build a two-layer hierarchical model with a support vector machine (SVM) (15). Subsequently, Sen *et al.* introduced PyPredT6, an integrated prediction tool that leverages nucleotide and amino acid sequence information of T6SEs, incorporating algorithms such as Multi-layer Perceptron (MLP), SVM, k-nearest neighbor (k-NN), Naive Bayes (NB), and Random Forest (RF) (19). Additionally, several comprehensive tools and algorithms have been developed to predict multiple secretion systems simultaneously. For instance, BastionX integrates Hidden Markov Models (HMMs) with machine learning techniques to predict T1SE-T4SE and T6SE independently (16). Tbooster employs three ensemble models that integrate various machine learning approaches to predict T3SE, T4SE, and T6SE (17), while Orgsissec uses phylogenetic features and sequence encoding for similar predictions (18).

Machine learning-based tools for T6SE prediction have become invaluable resources for identifying and characterizing T6SEs. However, traditional machine learning methods face limitations such as challenges in feature selection and susceptibility to data noise. Deep learning algorithms present a promising approach to address these limitations. Recently, deep learning-based prediction tools have demonstrated remarkable success in various domains, exemplified by T4SEpp and CNN-T4SE for effector prediction (20, 21). Moreover, current T6SE prediction tools heavily depend on training datasets with high sequence homology and limited positive samples, which can adversely affect model generalization. Thus, developing an integrated deep learning model for T6SE prediction with datasets of lower sequence homology is crucial. This approach would improve model performance by enhancing its ability to handle diverse sequences.

In this study, we introduce T6CNN, a novel ensemble approach utilizing convolutional neural networks (CNNs) for T6SE prediction. T6CNN integrates diverse protein features, such as N-terminal signal sequences and evolutionary data, to extract multi-level and multi-dimensional representations. These features are subsequently utilized to construct CNN-based classification models. T6CNN employs a voting strategy to identify potential T6SE candidates, combining predictions from individual CNN models to generate a final consensus. This ensemble strategy significantly improves the accuracy and robustness of T6SE prediction. The T6CNN tool is freely accessible at https://github.com/yuemhu/T6CNN, offering researchers a user-friendly platform for T6SE prediction using CNN-based algorithms.

## 2 Materials and methods

The model development and prediction (Figure 1) involved four major stages: Data collection and preprocessing, feature extraction, model construction and training, and performance evaluation. Each of these major stages is described in the following sections.

**Figure 1.**
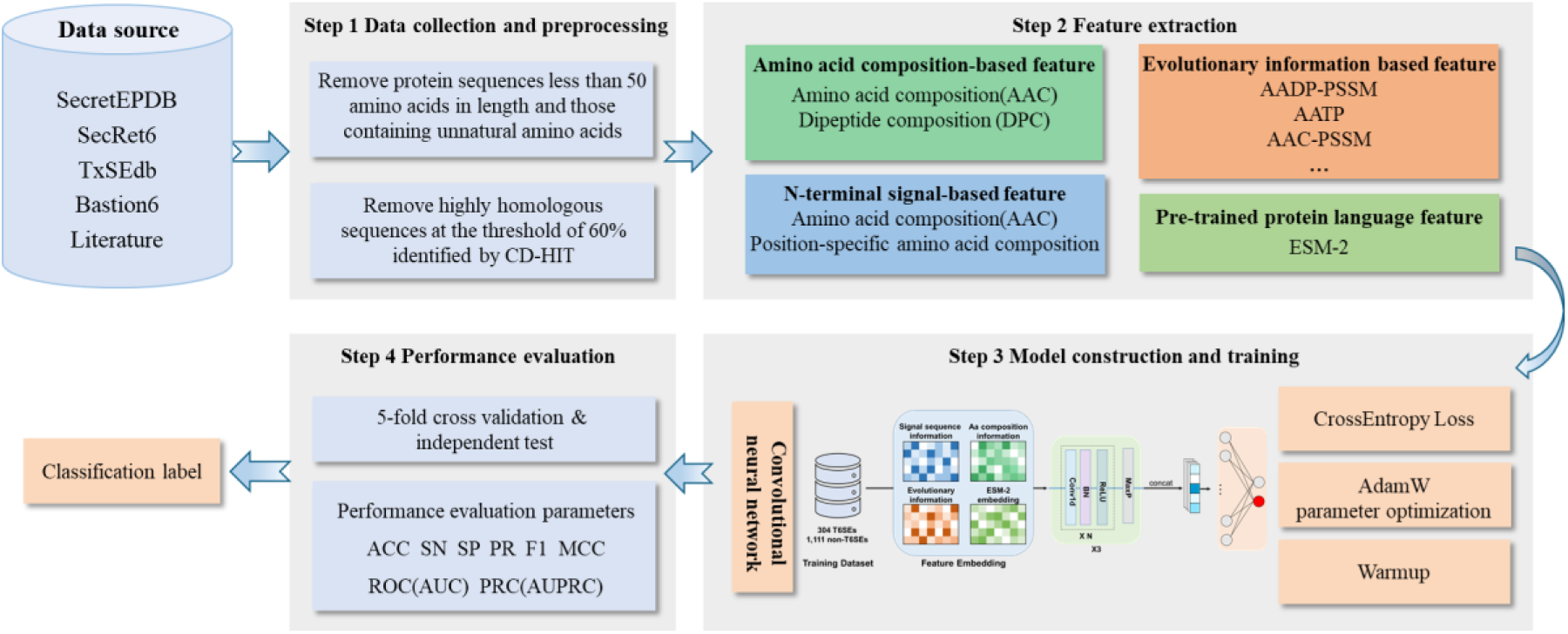
Overview of the proposed methodology for predicting T6SEs. First, a large dataset of verified T6SS effectors and non-effectors is collected from multiple databases and preprocessed to form clean training and test sets. Then, diverse features are extracted, including N-terminal signal descriptors, overall amino acid and dipeptide compositions, evolutionary profiles from PSSM, and contextual embeddings from the ESM-2 transformer model. Next, these heterogeneous features are integrated into the T6CNN model, which employs parallel convolutional layers, pooling, and feedforward networks with cross-entropy loss and AdamW optimization for training. Model performance is evaluated using 5-fold cross-validation and an independent test with various quantitative metrics. Finally, the well-trained model reliably distinguishes putative T6SEs from non-effectors.

### 2.1 Data collection and preprocessing

We collected and merged 138 T6SEs in Bastion6 (15), 181 T6SEs in SecretEPDB (22), 329 T6SEs in SecRet6 (23) and 111 T6SEs in TxSEdb (http://61.160.194.165:3080/TxSEdb/index.html) to get a total of 760 verified T6SEs. The same negative samples as Bastion6 were used, containing 1112 non-effector sequences. We excluded sequences less than 50 amino acids in length and those containing unnatural amino acids, and then removed homologous sequences by CD-HIT (24) at a threshold of 60% sequence identify. Finally, we obtained a non-redundant training dataset containing 305 positive and 1,111 negative protein sequences.

To generate the independent test set for evaluating the model performance in comparison with other existing T6SE prediction approaches, we combined 14 newly experimentally validated T6SEs from recent literature with 20 T6SEs from Bastion6 testing set (Supplementary Table S1). For negative samples, we utilized the same set as Bastion6, consisting of 200 non-T6SEs. Applying the same preprocessing as the training dataset, which involved removing highly homologous samples with over 60% similarity using CD-HIT, we obtained a non-redundant independent test set with 28 T6SEs and 185 non-T6SEs.

### 2.2 Feature extraction

The amino acid sequence of proteins carries intrinsic characteristics that can be transformed into numerical features for model inputs. Various feature descriptors capture diverse protein properties from different perspectives. In this study, we utilized four categories of feature descriptors, including N-terminal signal sequence amino acid composition-based features, amino acid composition-based features, evolutionary information-based features, and transformer protein language models (ESM-2)-based features. These descriptors contribute to the model performance and facilitate the interpretation of differences between T6SEs and non-T6SEs, offering valuable insights for further research.

#### 2.2.1 N-terminal signal sequence amino acid composition-based features

Bacterial secretion signal sequences have been widely acknowledged to play a critical role in directing effector proteins towards the secretion pathway. Various prediction tools based on signal sequences have also been developed to facilitate more accurate identification of effectors, such as T4SEpre, which utilizes the C-terminal sequential and position-specific amino acid compositions, possible motifs and structural features (25), and T3SEpp, which integrates promoter information along with characteristic protein features for signal regions (26). Although the investigation of T6SE signal sequences remains limited, Zhang et al. analyzed over 300 T6SEs using SignalP5.0 (27) and identified potential secretion signals in the N-terminal 30-70 amino acids of the majority of T6SEs (23). To further explore this, we extracted the N-terminal 50 and 100 amino acids from both T6SE and non-T6SE sequences, employing CD-HIT with a 30% threshold to eliminate highly similar fragments. By calculating the amino acid composition (Aac) within these regions, we derived a 20-dimensional frequency vector for each sequence, which was subsequently used to construct a frequency matrix.

The position-specific Aac features were extracted as follows. Suppose vector *S = s_1_*, *s_2_*, …, *s_i_*, …, *s_L_* denotes a protein sequence in which s represents an amino acid, *i* represents position and *L* represents sequence length. For *m* sequences, the position-specific Aac of a certain amino acid *A* is calculated by *p(A_i_) = f(A_i_)/m*, in which *f(A_i_)* represents the frequency of amino acid *A* at position *i*. For each position, the *p(A_i_)* of the 20 amino acid species can form a position-based set *M_i_*. Therefore, for the sequences of 50 and 100 amino acids in length, we can obtain 50-dimensional and 100-dimensional position-specific Aac frequency matrices by extracting corresponding elements from *M_i_*, respectively. We generated a 120-dimensional feature matrix by splicing the 20-dimensional Aac frequency matrix with the 100-dimensional position-specific Aac frequency matrix. Moreover, two 100-dimensional position-specific Aac frequency matrices extracted respectively from the *M_i_* of positive and negative proteins were spliced to a 200-dimensional feature matrix.

The amino acid composition and position-specific amino acid composition were compared between T6SEs and non-T6SEs with the Mann-Whitney U test, with a significance level set at p < 0.05. A log ratio of average frequency was then calculated for each amino acid species showing significant bias, representing their relative prevalence between positive and negative sequences. Similarly, for positions exhibiting significant bias, a log ratio of amino acid frequency was calculated across the 20 amino acid species, illustrating the distribution advantages at different positions. Sequence logos profiles were drawn to visualize the position-specific amino acid composition.

#### 2.2.2 Amino acid composition-based features

The primary structure of proteins provides functional information, which we captured through three types of amino acid composition-derived features, including amino acid composition (AAC) and dipeptide composition (DPC). AAC and DPC are feature descriptors used to capture different aspects of protein sequence information. AAC represents the occurrence frequencies of 20 amino acids, and DPC reflects sequence order information by considering the frequencies of dipeptides. These features were extracted using the iFeature online platform (28), resulting in 20-dimensional and 400-dimensional feature matrices, respectively.

#### 2.2.3 Evolutionary information-based features

Evolutionary conservation provides valuable insights into important biological functions beyond amino acid composition-based descriptors. Position-Specific Scoring Matrix (PSSM) is an effective feature encoding method that captures evolutionary information. It is generated using PSI-BLAST to search for homologous sequences and represents substitution scores between amino acid types (29). For each sequence, an *L*×20 matrix can be generated, in which *L* represents the length of a protein. The *(i, j)^th^* element of the matrix symbolizes the substitution score of the amino acid in the *i^th^*position mutating into amino acid type *j* during the evolutionary process. Based on PSSM, row and column transformations yield various feature descriptors. Row transformation-based descriptors include AAC-PSSM, S-FPSSM, AB-PSSM, PSSM-composition, RPM-PSSM, and smoothed-PSSM. Column transformation-based descriptors include DPC-PSSM, k-separated-bigrams-PSSM, tri-gram-PSSM, EEDP, and TPC-PSSM. Mixed row and column transformation-based descriptors include EDP, RPSSM, Pse-PSSM, DP-PSSM, PSSM-AC, and PSSM-CC. Combinative descriptors include AADP-PSSM, AATP, and MEDP. These descriptors range from 20-dimensional to 3800-dimensional matrices, capturing diverse evolutionary information. The detailed descriptions and calculation methods for the 20 feature descriptors can be found in the Supplementary Methods. Evolutionary information-based features were all extracted through the POSSUM online platform (30).

#### 2.2.4 Transformer protein language models (ESM-2)-based features

Transformer-based protein language models have shown great success in capturing universal representations of protein sequences. These models have been widely applied to various prediction tasks related to protein structure and properties. One such model, ESM-2, developed by Facebook AI Research, is a transformer-based protein language model with 650 million parameters, trained on the UniRef50 database (31). The model learns rich biological embeddings by processing diverse protein sequences, enabling it to capture intricate patterns of protein biology. In this study, we used the pre-trained ESM-2 model to extract protein sequence features without additional fine-tuning on our specific dataset. The model generates a feature matrix for each protein sequence, where the matrix has dimensions *L*×1280. Here, *L* represents the length of the protein sequence, and 1280 corresponds to the dimensionality of the embedding space provided by the ESM-2 model. The extracted embeddings, which serve as frozen representations of the proteins, encapsulate both local and global contextual information from the sequences.

### 2.3 Construction of T6CNN model

The T6CNN architecture used in this study is composed of three convolutional layers, a pooling layer, two feedforward neural networks (FNNs), and an output layer. Given an input sequence embedding z ∈ ℝ, then fed into three parallel convolutional layers, each applying convolution operations using three different convolution kernels with sizes *k_1_=n*, *k_2_=2n*, and *k_3_=3n*, where *n* is a hyperparameter. For the *n-th* convolution operation:

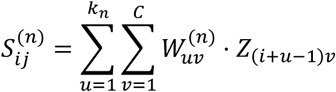

Here, *i* represents the position index in the output feature map along the sequence length, *j* represents each convolution channel index in the feature map, *u* represents the index along the convolution kernel height (length of the input sequence), *v* represents the input channel index, 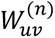 denotes the convolutional weights, and *Z*_(*i*+*u*−1)*v*_ is the input value at the corresponding location. Each convolutional output is followed by a batch normalization, rectified linear unit (ReLU) activation, and dropout operation:

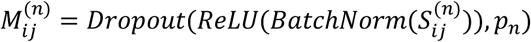

The pooling layer employs max-pooling to select the maximum neuron value over a region, reducing variance and increasing translational invariance. The pooled feature maps from the three convolutional layers are concatenated into a single vector:

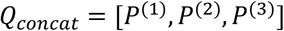

This vector is then passed through FNNs. Finally, the output layer applies a linear transformation followed by a softmax activation function to generate the classification probabilities.

### 2.4 Training of T6CNN model

In this study, the cross-entropy loss function was utilized for training the models, and the AdamW optimizer was employed to adjust the parameters with adaptive learning rates. The initial learning rate and weight decay were set to 0.001 and 0.025, respectively. To optimize the learning rate, the cosine Warmup strategy with 20 iterations was implemented. To address potential overfitting, a dropout strategy was applied with a dropout probability of 0.25, randomly setting input elements to zero during training. T6CNN was developed with PyTorch v1.10.1. The models were trained and evaluated with 24-GB memory and NVIDIA GeForce RTX 3090 GPU for acceleration.

### 2.5 Performance evaluation

To evaluate the prediction performance, we first used principal component analysis (PCA) to carry out dimensionality reduction for each feature, which was visualized as a two-dimensional projection of the embedding spaces using t-SNE. Combined with the K-means algorithm, the performance of each feature coding method was preliminarily evaluated by unsupervised clustering. Then, we further performed the supervised analysis, which included 5-fold cross-validation and independent test. Six quantification metrics, including Accuracy (ACC), Sensitivity (SN), Specificity (SP), Precision (PR), F1-score (F1), and Matthew’s correlation coefficient (MCC) were used to evaluate the overall predictive performance of classification models. These are defined as follows:

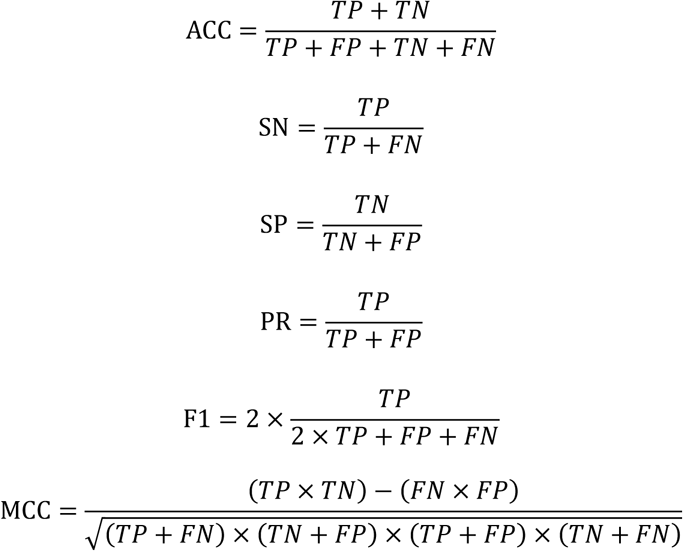

where TP, TN, FP and FN denote the numbers of true positives, true negatives, false positives and false negatives, respectively.

Additionally, the receiver operating characteristic (ROC) curve, which is a plot of the true-positive rate versus the false-positive rate, and the precision-recall curve (PRC), which is a plot of the precision versus the recall, are depicted to visually measure the comprehensive performance of different feature coding methods. The area under the curves termed AUC and AUPRC is also provided in each of the plots. For imbalanced data, the precision-recall curve is a more suitable choice than the ROC curve.

## 3 Results

### 3.1 Sequence-based amino acid composition (Aac) differences between N-terminal of T6SEs and non-T6SEs

Based on previous studies, we hypothesized that the T6SE signal sequence is located within the 30-70 amino acid (aa) region at the N-terminus (23, 27). Therefore, we compared the sequence-based Aac of the first 50 and 100 positions in the N-terminal regions of T6SEs and non-T6SEs. The distributions of most amino acids differed significantly between the two sequence types. Alanine, cysteine, aspartic acid, glycine, proline, serine, threonine, tryptophan, and tyrosine were enriched, while phenylalanine, isoleucine, lysine, leucine, methionine, arginine, and valine were depleted in T6SEs (Mann-Whitney U test, p<0.05, Figure 2). In both the 50-aa (Figure 2A) and 100-aa (Figure 2B) fragments, cysteine, aspartic acid, glycine, and serine were enriched, while phenylalanine, isoleucine, leucine, and methionine were depleted.

**Figure 2.**
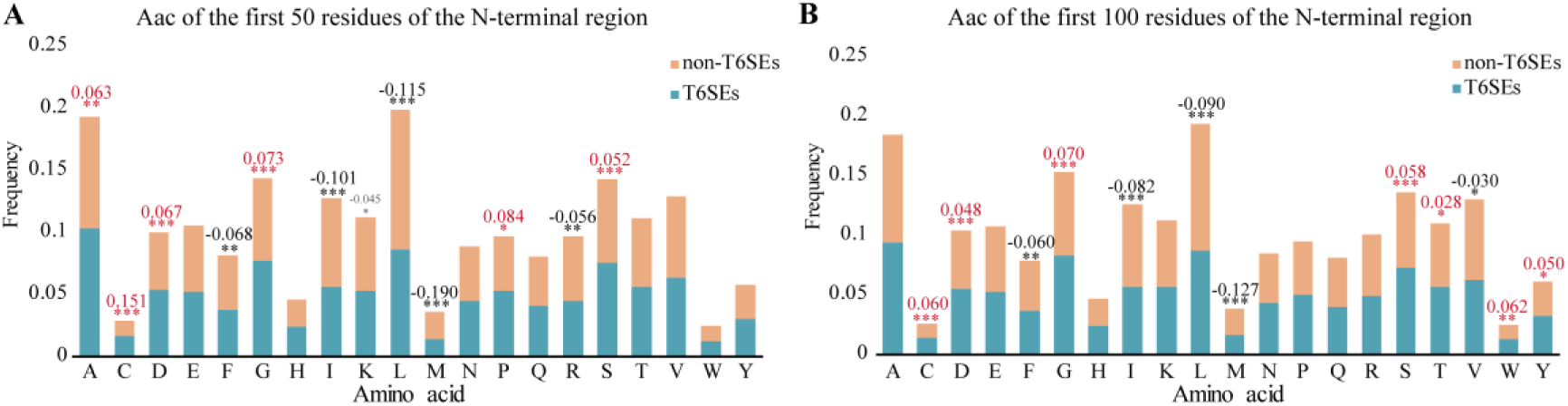
Comparison of amino acid composition in the N-terminal regions of T6SEs and non-T6SEs. (A) Amino acid composition of the first 50 residues of the N-terminal region in T6SEs and non-T6SEs. (B) Amino acid composition of the first 100 residues of the N-terminal region in T6SEs and non-T6SEs. The x-axis lists the 20 standard amino acids, with bar lengths representing their relative frequencies. T6SEs and non-T6SEs are represented by blue and orange bars, respectively. Amino acids that show significant differences (Mann-Whitney U test, p < 0.05 *; p < 0.01 **; p < 0.001 ***) are marked with asterisks above the bars. Additionally, the logarithmic ratio of average amino acid frequency between T6SEs and non-T6SEs is shown, with red indicating enrichment in T6SEs and black indicating depletion.

### 3.2 Position-specific amino acid composition differences between N-terminal of T6SEs and non-T6SEs

We compared the position-specific Aac profiles of T6SEs and non-T6SEs, revealing clear differences. T6SEs exhibited distinct amino acid preferences at most positions, whereas non-T6SEs showed a more uniform amino acid distribution (Figure 3, Supplementary Figure S1). A comparison of position-specific Aac in the first 100 N-terminal positions (excluding position 1, predominantly methionine) showed that approximately 40% of positions exhibited significant differences (Supplementary Table S2). These differences were distributed across the sequence, with a slight concentration around positions 46-53.

**Figure 3.**
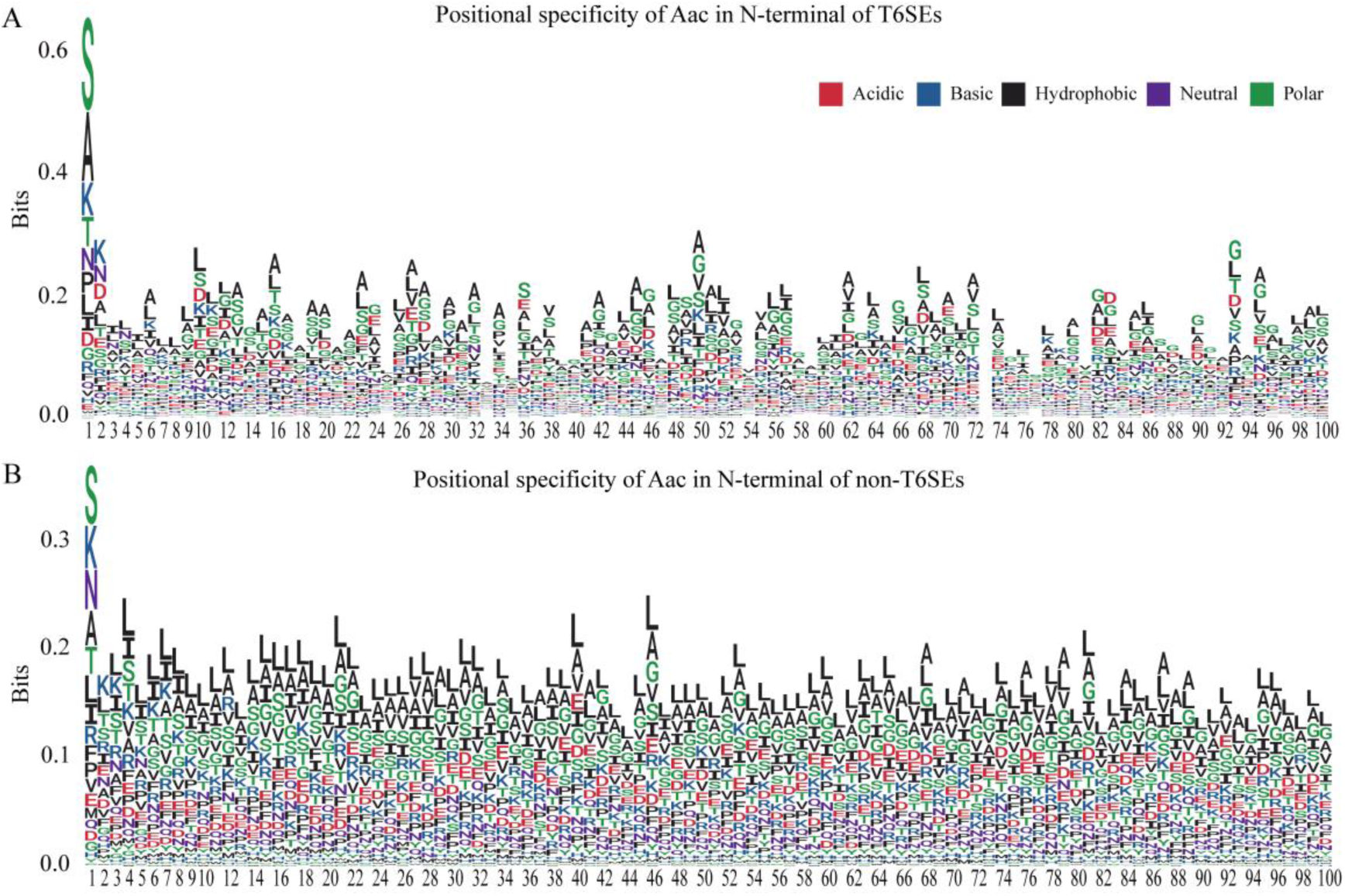
Positional specificity of amino acid composition in the N-terminal regions of T6SEs and non-T6SEs. (A) Positional specificity of amino acid composition in the first 100 residues of the N-terminal region of T6SEs. (B) Positional specificity of amino acid composition in the first 100 residues of the N-terminal region of non-T6SEs. The x-axis represents the position in the N-terminal sequence, while the y-axis indicates the information content at each position in bits. The first position, typically occupied by methionine (M) in most proteins, is excluded. The stacked height at each position reflects the level of conservation, with the size of the letters representing the relative frequency of each amino acid.

For positions with significant differences (p<0.01; N1, N10, N13, N27, N36, N43, N47, N49, N50, N52, N56, N68, N72, N83, N93, and N95), we analyzed the distribution preferences of all 20 amino acids. As shown in Figure 4, the most pronounced differences were observed in the frequencies of cysteine, methionine, histidine, and tryptophan between T6SEs and non-T6SEs. Cysteine was sparsely distributed across most T6SE positions (N13, N50, N68, N83, and N93) but was enriched at position 47. Histidine and methionine were generally less frequent in T6SEs than in non-T6SEs. Tryptophan was significantly enriched at positions 47, 56, and 83, but depleted at positions 10, 27, 36, 49, and 50 in T6SEs.

**Figure 4.**
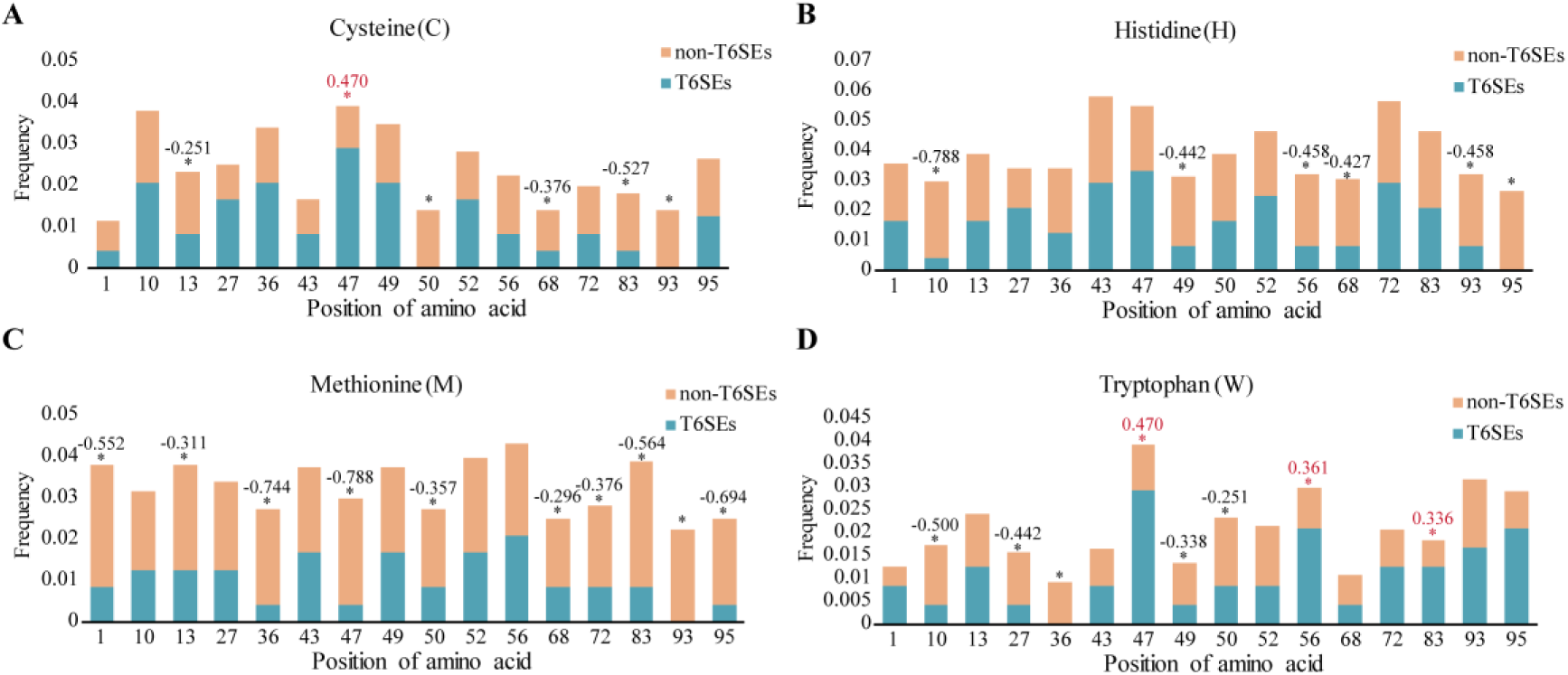
Differential distribution of specific amino acids at specific positions in the N-terminal regions of T6SEs and non-T6SEs. (A) Distribution of cysteine at specific positions in the N-terminal region of T6SEs and non-T6SEs. (B) Distribution of histidine at specific positions. (C) Distribution of methionine at specific positions. (D) Distribution of tryptophan at specific positions. The x-axis lists sixteen sequence positions, with bar lengths representing the amino acid frequency at each position. T6SEs and non-T6SEs are represented by blue and orange bars, respectively. Significant differences in distribution are marked with asterisks above the bars. Additionally, the logarithmic ratio of amino acid frequencies is shown, with red text indicating relative enrichment in T6SEs and black text indicating relative depletion in T6SEs.

The enrichment or depletion of specific amino acids in T6SE sequences may relate to secondary structure and hydrophilicity, factors potentially critical for signal recognition. Negatively charged aspartic acid and glutamic acid at the N-terminus reportedly influence protein secretion, while interactions between positively charged histidine and negatively charged residues may also affect secretion. However, further investigation is needed to fully understand the biological mechanisms underlying these amino acid biases. Overall, the findings suggest that the features extracted from N-terminal sequences serve as an effective coding strategy for distinguishing T6SEs from non-T6SEs.

### 3.3 Unsupervised Analysis of Feature Encoding Methods for T6SE Classification

To intuitively estimate the effect of different feature encoding methods on the classification performance, we adopted principal component analysis (PCA) and t-distributed stochastic neighbor embedding (t-SNE) to map all the samples onto two-dimensional space. As shown in Figure 5, negative samples were evenly distributed across the two-dimensional map, indicating that they were representative. Most evolutionary information-based features including AAC-PSSM, AB-PSSM, DP-PSSM, EEDP, MEDP, Pse-PSSM, PSSM-AC, PSSM-CC, PSSM-composition, RPM-PSSM, RPSSM, and amino acid composition-based features, including AAC, performed well, under which most positive samples could be clustered together (Figure 5, Supplementary Figure S2). However, for N-terminal signal sequence amino acid composition-based features, both types of samples showed scattered distribution (Figure 5A, Supplementary Figure S2A).

**Figure 5.**
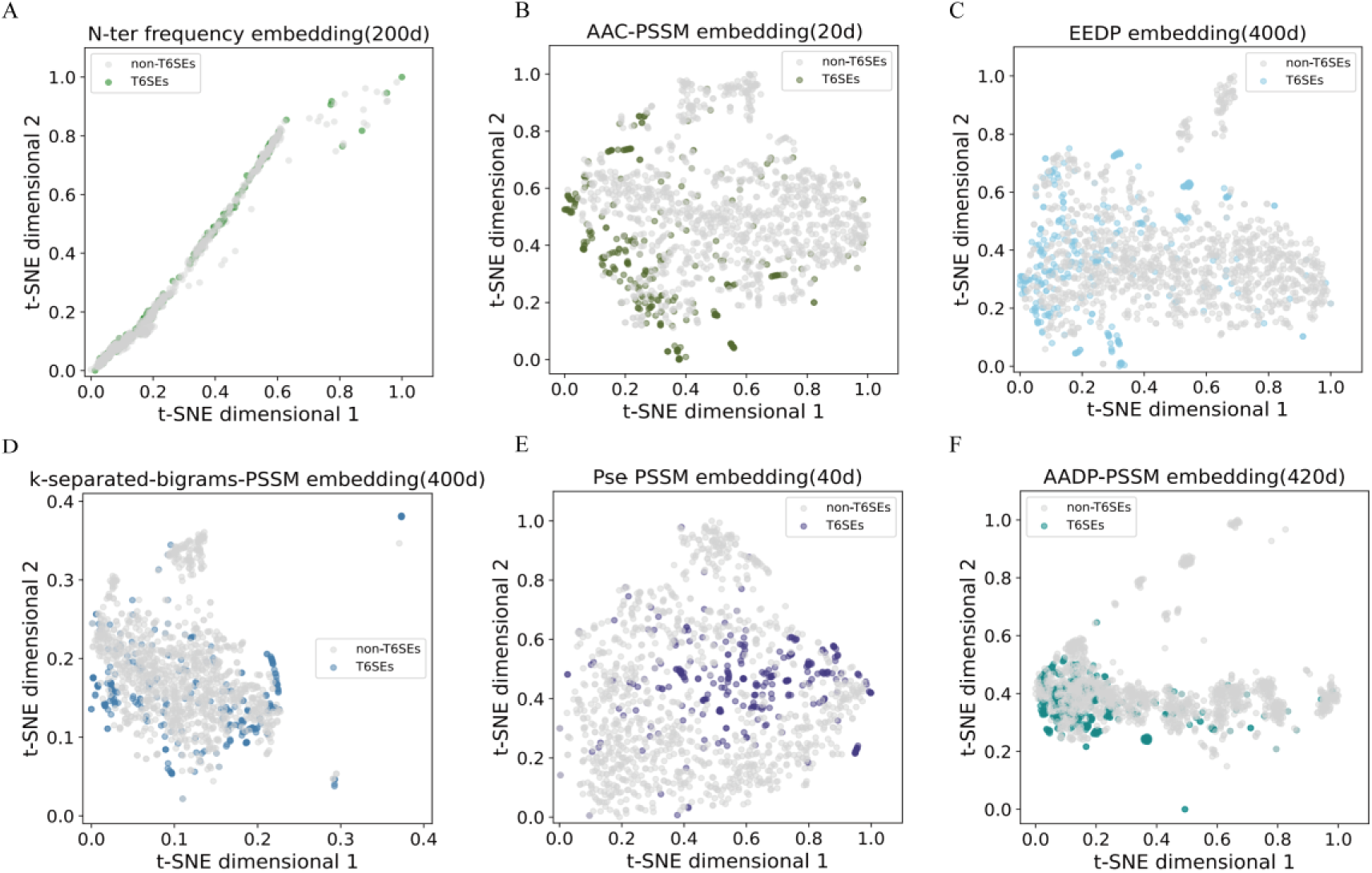
t-SNE visualization of embedding vectors generated by different feature encoding strategies. (A-F) show the distribution of positive (colored) and negative (gray) samples after PCA dimensionality reduction, with each point representing a sample. The color coding indicates the distribution of positive samples under different feature encoding strategies.

To address the lack of clear classification in the embedding, K-means clustering was applied to the data. RPM-PSSM outperformed other feature encodings, under which non-T6SEs were mostly distributed in Cluster 1 (accounting for 61.4%) while T6SEs dominated in Cluster 2 (accounting for 86.2%, Supplementary Figure S3, Supplementary Table S2). In general, evolutionary information-based features contributed more to T6SEs classification, while all feature encoding methods showed high mixing rates within each cluster, which further illustrated the necessity of constructing a classification model using supervised analysis.

### 3.4 Supervised analysis

To evaluate the performance of each feature encoding method, a T6CNN model was trained and assessed using 5-fold cross-validation and an independent test.

#### 3.4.1 Performance evaluation using 5-fold cross-validation

In 5-fold cross-validation, the samples were randomly divided into five equal sized subsets, with four subsets used for training data and one subset for testing. This process was repeated five times, recording the performance metrics each time and calculating the average performance. As illustrated in Supplementary Table S3 and Figure 6, evolutionary information-based features, such as AAC-PSSM, AADP-PSSM, AATP, EEDP, DPC-PSSM, k-separated-bigrams-PSSM, Pse-PSSM and PSSM-composition, demonstrated overall good performance with ACC (>0.91), SN (>0.75), SP (>0.94), PR (>0.81), F1 (>0.80) and MCC (>0.75). Among them, EEDP and AATP displayed a better performance in terms of SP (0.969±0.013) and PR (0.877±0.062), respectively. The performance of AAC- and DPC-based feature was inferior to the feature encoding methods based on evolutionary information. These findings reaffirm that PSSM-based features provided more informative representations compared to features based solely on amino acid composition.

**Figure 6.**
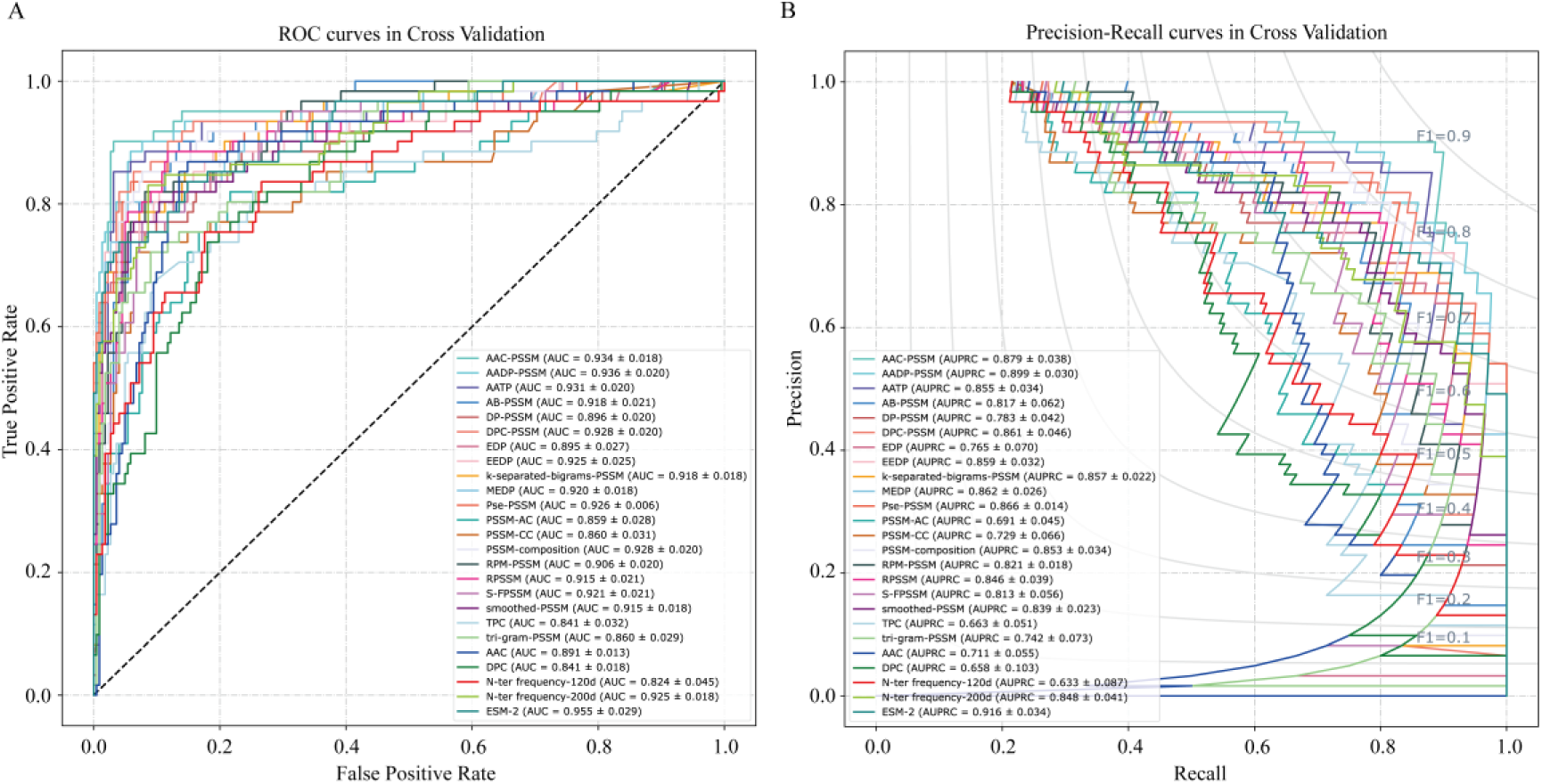
ROC and PRC curves for 25 feature encoding methods and their corresponding T6CNN models in 5-fold cross-validation. (A) ROC curves for 25 feature encoding methods and their corresponding T6CNN models in 5-fold cross-validation. The black dashed line represents the ROC curve for random label prediction. (B) PRC curves for 25 feature encoding methods and their corresponding T6CNN models in 5-fold cross-validation. Different feature encoding methods are represented by different colors. The area under the curve (AUC and AUPRC) is reported as mean ± standard deviation.

In terms of features based on N-terminal signal sequence amino acid composition, the 200-dimensional feature matrix demonstrated superior performance, as indicated by the ACC (0.905±0.021), SN (0.799±0.057), SP (0.934±0.028), PR (0.772±0.075), F1 (0.782±0.042), MCC (0.724±0.054), AUC (0.925±0.018) and AUPRC (0.848±0.041) (Figure 6, Supplementary Table S3). ESM-2-based features emerged as the optimal feature encoding strategy, achieving the highest average values for ACC (0.939±0.012), SN (0.845±0.061), F1 (0.855±0.027), and MCC (0.818±0.033). Additionally, ESM-2-based feature exhibited the best performance in terms of AUC (0.955±0.029) and AUPRC (0.916±0.034).

Compared to unsupervised analysis, certain feature encoding methods such as AAC-PSSM, Pse-PSSM、PSSM-composition, and EEDP consistently demonstrated excellent performance in both PCA and 5-fold cross-validation (Supplementary Figure S2, Supplementary Table S3). However, the results of AADP-PSSM, AATP, DPC-PSSM, k-separated-bigrams-PSSM, and N-terminal 200-dimensional frequency matrix showed discrepancies between PCA and 5-fold cross-validation, performing better in the latter. This indicates that the classification algorithm has a significant impact on the performance of the feature encoding methods.

#### 3.4.2 Performance validation on the independent test

The proposed models were further assessed using an independent test. As shown in Figure 7 and Supplementary Table S4, ESM-2 outperformed all other feature encoding methods, as revealed by the ACC (0.944), SN (0.966), SP (0.941), PR (0.718), F1 (0.824), MCC (0.803), and AUPRC (0.940). Additionally, N-terminal 200-dimensional frequency matrix displayed the second-best performance in terms of SN (0.964), F1 (0.761), MCC (0.739), and EEDP achieved an equivalent performance reflected by ACC (0.925), SP (0.930), PR (0.658). Following them, AAC-PSSM, k-separated-bigrams-PSSM, MEDP, RPSSM and Pse-PSSM also achieved satisfactory level in the case of ACC>0.90, SN>0.89, SP>0.90, PR>0.59, F1>0.72, MCC>0.69.

**Figure 7.**
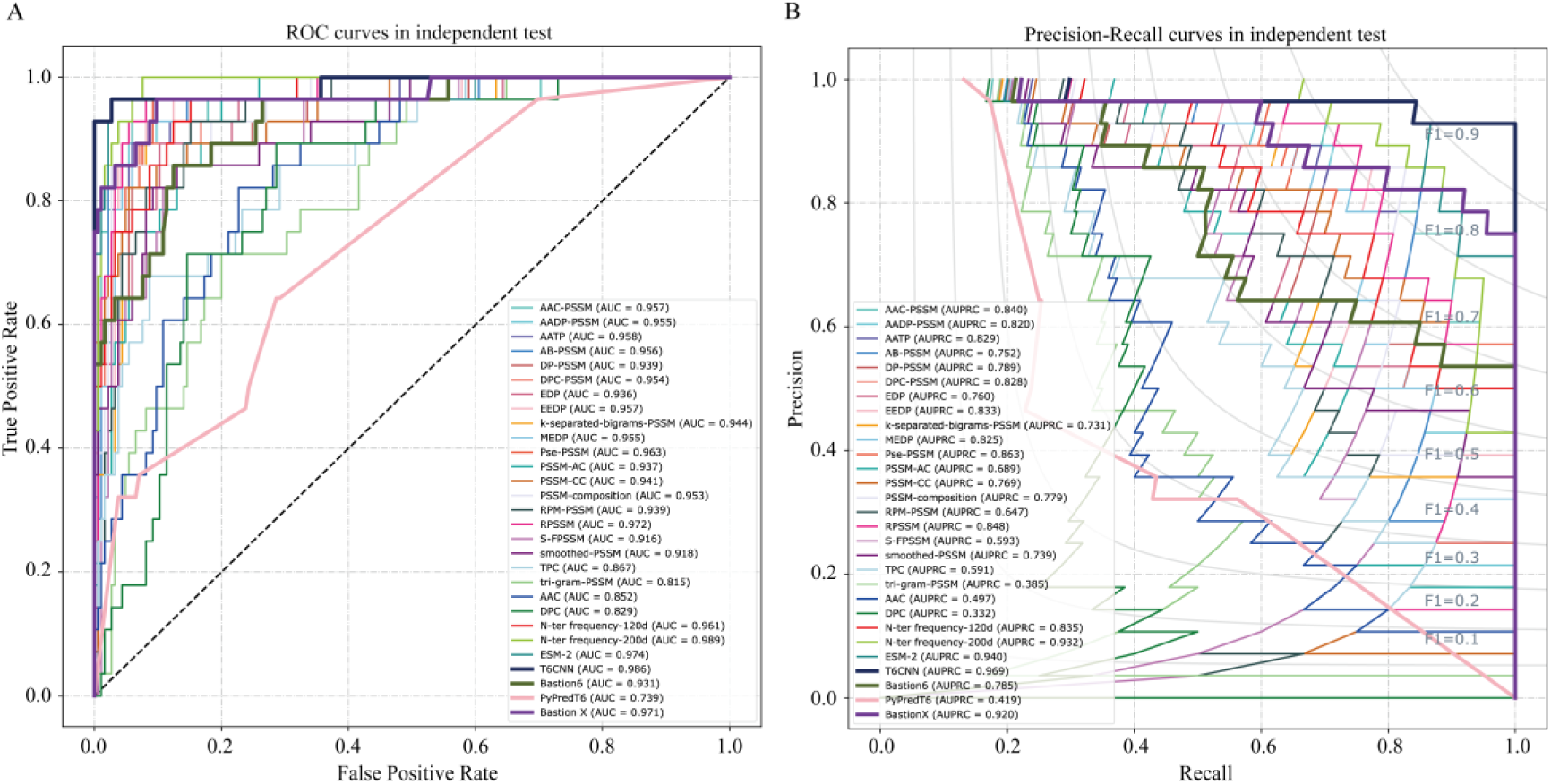
ROC and PRC curves comparing T6CNN with existing prediction tools in independent testing. (A) ROC curves and (B) PRC curves showing the performance of 25 different feature encoding methods, our ensemble model, and previous SOTA tools (Bastion6, PyPredT6, and BastionX). The black dashed line in panel A represents random prediction. Different methods are distinguished by unique colors.

In general, an ideal model had to strike a balance between SN and SP to achieve overall excellent performance, as they cannot both reach the highest values simultaneously. In our independent test, the gaps between SN and SP were minimal across all the feature encoding methods, indicating that our models demonstrated stable performances. After comprehensive evaluation of all the feature encoding methods based on unsupervised analysis, 5-fold cross-validation, and independent test, we constructed an ensemble model that integrated AAC-PSSM, EEDP, k-separated-bigrams-PSSM, Pse-PSSM, AADP-PSSM, and N-terminal 200-dimensional frequency matrix and ESM-2, along with their corresponding T6CNN. The final class label was determined by a voting strategy, where a candidate was classified as a T6SE if at least three classifiers predicted it as a positive protein; otherwise, it was considered a non-T6SE. The proposed ensemble classifier T6CNN performed well on the independent test, as demonstrated in Figure 7 and Table 1, achieving high values for ACC (0.953), SN (0.964), SP (0.951), PR (0.750), F1 (0.844), MCC (0.826), AUC (0.986) and AUPRC (0.969).

**Table 1.** Performance of the existing methods and our methods.

| Methods | ACC | SN | SP | PR | F1 | MCC |
| --- | --- | --- | --- | --- | --- | --- |
| Bastion6 | 0.883 | 0.821 | 0.892 | 0.535 | 0.648 | 0.600 |
| PyPredT6 | 0.718 | 0.500 | 0.751 | 0.233 | 0.318 | 0.189 |
| Bastion X | 0.859 | **0.964** | 0.843 | 0.482 | 0.643 | 0.620 |
| Our Method | **0.953** | **0.964** | **0.951** | **0.750** | **0.844** | **0.826** |

Note: ACC: accuracy; SN: sensitivity; SP: specificity; PR: precision; F1: F1-score; MCC: Matthews correlation coefficient. For each metric, the best performance value across different encoding methods is highlighted in bold.

### 3.5 Performance comparison with existing tools

In comparison to existing bioinformatics tools for T6SE prediction, such as Bastion6, PyPredT6, and BastionX, T6CNN outperforms them in independent tests across all 8 performance indicators. Our method achieves the highest scores in terms of accuracy, sensitivity, and specificity, with an accuracy of 0.953 (improved by 7.9%), sensitivity of 0.964, specificity of 0.951 (increased by 6.6%), precision of 0.750 (exceeding by 40.2%), F1-score of 0.844 (improved by 30.2%), Matthews’ correlation coefficient of 0.826 (increased by 37.7%) (Figure 7, Table 1). This demonstrates the superior performance of our method compared to the previous state-of-the-art (SOTA) tool.

## Discussion

Gram-negative bacteria encode T6SS to achieve bacterial interaction, antibacterial competition, and virulence to eukaryotic hosts, thereby improving the adaptability of bacteria to the environment and enhancing their own viability (32, 33). Understanding the functioning of T6SS in these interactions is crucial. Identifying potential T6SEs is a key step in this process, and bioinformatics methods, ranging from simple alignments to machine learning, have been developed. However, these methods have limitations due to the small number of known T6SEs and species-specific motifs. To address these challenges and enable accurate T6SE prediction across diverse bacterial species, we propose a novel genome-scale effector protein prediction method based on convolutional neural networks.

In this study, we developed T6SE prediction tools based on protein sequence features, focusing on the analysis of the N-terminal signal sequence. We found that the amino acid composition of the N-terminal sequence has a distinct preference in T6SEs compared to non-T6SEs. We extracted a total of 25 features to capture protein sequence information from multiple perspectives, including N-terminal sequence composition and evolutionary information. Different features showed varying performance in different analysis methods, with PSSM-based features performing well overall. Simple features based solely on amino acid composition performed poorly. The N-terminal sequence composition showed promising results in cross-validation and independent testing. We selected six single-feature models and employed a voting strategy to improve the prediction performance of the final model for T6SEs. By leveraging different features and combining them appropriately, we enhanced the accuracy of T6SE prediction.

Compared to existing T6SE prediction methods, our approach demonstrates superior performance across multiple evaluation parameters, including accuracy, sensitivity, specificity, precision, F1 score, and MCC (15, 16). Our research offers several advantages and innovations. Firstly, we collected a more comprehensive set of experimentally verified T6SEs, increasing the size of the positive sample set. Secondly, we implemented a lower threshold (60%) for filtering homology redundancy compared to previous methods, which used a threshold of 90%. This reduced sequence redundancy and improved the generalization of our model. Thirdly, we introduced the novel application of extracting amino acid composition features from the N-terminal signal sequence, achieving excellent prediction performance. Fourthly, our utilization of a deep learning algorithm provides advantages in terms of prediction speed, model generalization, and reduced sensitivity to noise. However, our method has certain limitations, such as the inability to process sequences shorter than 50 amino acids or those containing unnatural amino acids. Additionally, we solely relied on protein sequence features, excluding other known features that aid in T6SEs identification. Integration of features like GC content, phylogenetic maps, regulatory motifs, and physicochemical properties could enhance the model’s performance (6). Future research should focus on uncovering novel features to further improve the predictive capabilities of our model.

Therefore, our method demonstrates outstanding performance and holds immense potential as a valuable tool for T6SE prediction. Its accuracy and efficiency can greatly expedite the discovery of new T6SEs, deepening our understanding of bacterial T6SS functionality. This has far-reaching implications, particularly in the development of novel antibacterial drugs. By facilitating the identification of T6SEs, our method significantly advances research in this area and enables targeted interventions against bacterial pathogens. It opens up promising avenues for further exploration and investigation in the field of T6SE prediction, fostering advancements in our knowledge of bacterial-host interactions.

## Data availability

The standalone version of T6CNN model and the individual modules are also deposited at https://github.com/yuemhu/T6CNN.

## Acknowledgments

This work was supported by the National Key Research and Development Program of China (2023YFE0112300); the National Natural Sciences Foundation of China (32270709, 32261133526, 32300532); the Science and Technology Innovation Leading Scientist (2022R52035) and the 151 talent project of Zhejiang Province (first level); Postdoctoral Fellowship Program of CPSF (GZC20232322); Collaborative Innovation Center for Modern Crop Production co-sponsored by province and ministry.

## Authors’ Contribution

**Project Conception and Supervision:** M.C., Y.H., and C.F. conceived and supervised the project.

**Project Coordination:** Y.H. and M.Y. coordinated the project.

**Data Collection:** Y.H., M.Y., Y.Z., and Y.W. were responsible for dataset collection.

**Code, Models, and Software Tools:** Y.H. and M.Y. provided the codes, models, and software tools.

**Figure Creation:** Y.H., M.Y., E.L., and L.L. created the figures.

**Writing of the First Draft:** Y.H., M.Y., H.C., S.L., Q.N., YS.H., E.L., L.L., Y.C., Z.Z., Y.W., C.F., and M.C. wrote the first draft of the manuscript.

**Manuscript Revision:** Y.H., M.Y., Y.C., S.Z., Y.W., C.F., and M.C. revised the manuscript.

## Conflict of interest

The authors declare no conflict of interest.

---

**Supplementary Table S1:** Experimentally Validated Type VI Secretion System Effectors (T6SEs) Identified in Recent Literature (June 2020–December 2022)

| Protein ID | Species | Reference (DOI) |
| --- | --- | --- |
| CCC31770.1 | *Salmonella bongori* | 10.7554/eLife.82437 |
| CCC31775.1 | *Salmonella bongori* | 10.7554/eLife.82437 |
| CCC30909.1 | *Salmonella bongori* | 10.7554/eLife.82437 |
| CCC30901.1 | *Salmonella bongori* | 10.7554/eLife.82437 |
| WP_021703699.1 | *Vibrio proteolyticus* | 10.7554/eLife.82766 |
| WP_021703698.1 | *Vibrio proteolyticus* | 10.7554/eLife.82766 |
| WP_064704461.1 | *Pantoea agglomerans* | 10.1111/1462-2920.16100 |
| QUU06308.1 | *Bacteroides caccae* | 10.1128/jb.00122-22 |
| QUT66662.1 | *Bacteroides uniformis* | 10.1128/jb.00122-22 |
| QUT66677.1 | *Bacteroides uniformis* | 10.1128/jb.00122-22 |
| QUU06293.1 | *Bacteroides caccae* | 10.1128/jb.00122-22 |
| BAC59653.1 | *Vibrio parahaemolyticus* | 10.15252/embr.202153981 |
| 7XUX_A/B | *Y. pseudotuberculosis* | 10.1038/s41467-022-35522-9 |
| ACY86862.1 | *Salmonella serovars* | 10.1016/j.celrep.2020.107813 |

**Supplementary Table S2.** Statistical analysis of position-specific amino acid composition frequency in the N-terminal 100 positions of T6SEs and non-T6SEs.

| Mann Whitney U test |  |  |  |  |  |  |  |  |  |
| --- | --- | --- | --- | --- | --- | --- | --- | --- | --- |
| Position | P value | Position | P value | Position | P value | Position | P value | Position | P value |
| 1 | **1.53e-8** | 21 | 0.2550 | 41 | 0.3209 | 61 | **0.0318** | 81 | 0.1504 |
| 2 | **0.0299** | 22 | **0.0144** | 42 | 0.3983 | 62 | 0.2792 | 82 | **0.0129** |
| 3 | 0.9413 | 23 | **0.0200** | 43 | **0.0056** | 63 | 0.0714 | 83 | **0.0026** |
| 4 | 0.9041 | 24 | **0.0226** | 44 | 0.0675 | 64 | **0.0156** | 84 | 0.0571 |
| 5 | 0.8354 | 25 | 0.3036 | 45 | 0.0780 | 65 | 0.3500 | 85 | 0.4585 |
| 6 | **0.0166** | 26 | 0.3141 | 46 | **0.0328** | 66 | **0.0349** | 86 | 0.3937 |
| 7 | 0.2781 | 27 | **0.0005** | 47 | **0.0007** | 67 | **0.0183** | 87 | 0.7733 |
| 8 | 0.9180 | 28 | 0.1464 | 48 | **0.0270** | 68 | **0.0093** | 88 | 0.7653 |
| 9 | 0.1986 | 29 | 0.6688 | 49 | **0.0072** | 69 | 0.0861 | 89 | 0.7319 |
| 10 | **0.0024** | 30 | 0.0552 | 50 | **0.0012** | 70 | 0.1417 | 90 | **0.0209** |
| 11 | 0.0613 | 31 | 0.7962 | 51 | **0.0169** | 71 | 0.1660 | 91 | 0.0572 |
| 12 | 0.1871 | 32 | **0.0291** | 52 | **0.0003** | 72 | **0.0015** | 92 | 0.0889 |
| 13 | **0.0002** | 33 | 0.1588 | 53 | **0.0422** | 73 | 0.4005 | 93 | **0.0022** |
| 14 | **0.0143** | 34 | 0.0605 | 54 | 0.1800 | 74 | 0.0712 | 94 | **0.0117** |
| 15 | 0.1402 | 35 | 0.4308 | 55 | 0.0921 | 75 | 0.9800 | 95 | **0.0075** |
| 16 | 0.1480 | 36 | **0.0018** | 56 | **0.0033** | 76 | 0.3109 | 96 | 0.3777 |
| 17 | 0.1234 | 37 | 0.1750 | 57 | **0.0447** | 77 | 0.2288 | 97 | 0.6923 |
| 18 | 0.5010 | 38 | **0.0334** | 58 | 0.1119 | 78 | 0.4200 | 98 | 0.0539 |
| 19 | **0.0185** | 39 | 0.6855 | 59 | 0.8831 | 79 | 0.3267 | 99 | 0.1658 |
| 20 | **0.0189** | 40 | 0.8159 | 60 | 0.8648 | 80 | 0.0945 | 100 | **0.0481** |

Note: p<0.05 is highlighted in bold

**Supplementary Table S3.** Performance of various classifiers in this study evaluated by the 5-fold cross-validation.

| Encoding | ACC | SN | SP | PR | F1 | MCC |
| --- | --- | --- | --- | --- | --- | --- |
| **PSSM-based feature** |  |  |  |  |  |  |
| AAC-PSSM | 0.924±0.030 | 0.839±0.041 | 0.947±0.038 | 0.824±0.102 | 0.828±0.057 | 0.782±0.075 |
| AADP-PSSM | 0.934±0.016 | 0.826±0.072 | 0.964±0.018 | 0.867±0.055 | 0.843±0.041 | 0.805±0.047 |
| AATP | 0.933±0.016 | 0.806±0.068 | 0.968±0.019 | **0.877±0.062** | 0.833±0.046 | 0.799±0.049 |
| AB-PSSM | 0.903±0.022 | 0.774±0.027 | 0.939±0.026 | 0.782±0.078 | 0.776±0.043 | 0.716±0.058 |
| DP-PSSM | 0.898±0.011 | 0.744±0.025 | 0.940±0.017 | 0.775±0.047 | 0.758±0.019 | 0.695±0.027 |
| EDP | 0.888±0.029 | 0.744±0.030 | 0.928±0.032 | 0.746±0.082 | 0.744±0.053 | 0.674±0.071 |
| EEDP | 0.924±0.006 | 0.760±0.055 | **0.969±0.013** | 0.876±0.041 | 0.812±0.020 | 0.769±0.021 |
| DPC-PSSM | 0.931±0.020 | 0.797±0.056 | 0.967±0.019 | 0.874±0.067 | 0.832±0.049 | 0.791±0.062 |
| k-separated-bigrams-PSSM | 0.919±0.017 | 0.777±0.052 | 0.958±0.025 | 0.843±0.082 | 0.805±0.034 | 0.757±0.047 |
| MEDP | 0.917±0.020 | 0.761±0.034 | 0.960±0.018 | 0.843±0.066 | 0.799±0.044 | 0.750±0.058 |
| Pse-PSSM | 0.916±0.017 | 0.800±0.018 | 0.948±0.023 | 0.813±0.070 | 0.805±0.034 | 0.752±0.047 |
| PSSM-AC | 0.835±0.031 | 0.800±0.061 | 0.845±0.046 | 0.594±0.065 | 0.678±0.041 | 0.586±0.054 |
| RPSSM | 0.915±0.018 | 0.777±0.025 | 0.952±0.021 | 0.821±0.064 | 0.797±0.035 | 0.744±0.048 |
| PSSM-composition | 0.918±0.004 | 0.777±0.047 | 0.956±0.010 | 0.830±0.026 | 0.802±0.016 | 0.751±0.016 |
| RPM-PSSM | 0.912±0.007 | 0.748±0.038 | 0.958±0.007 | 0.830±0.019 | 0.786±0.020 | 0.733±0.023 |
| S-FPSSM | 0.903±0.015 | 0.774±0.050 | 0.939±0.010 | 0.777±0.033 | 0.775±0.036 | 0.713±0.045 |
| smoothed-PSSM | 0.908±0.017 | 0.751±0.048 | 0.951±0.023 | 0.815±0.071 | 0.779±0.036 | 0.724±0.049 |
| TPC | 0.833±0.031 | 0.715±0.015 | 0.866±0.042 | 0.603±0.080 | 0.651±0.042 | 0.549±0.058 |
| tri-gram-PSSM | 0.862±0.053 | 0.744±0.044 | 0.895±0.078 | 0.693±0.150 | 0.707±0.070 | 0.628±0.097 |
| PSSM-CC | 0.866±0.040 | 0.757±0.027 | 0.896±0.052 | 0.682±0.112 | 0.714±0.063 | 0.633±0.085 |
| **Amino acid composition-based feature** |  |  |  |  |  |  |
| AAC | 0.853±0.027 | 0.770±0.064 | 0.876±0.043 | 0.638±0.064 | 0.694±0.036 | 0.607±0.047 |
| DPC | 0.830±0.021 | 0.761±0.060 | 0.850±0.034 | 0.586±0.052 | 0.659±0.030 | 0.560±0.042 |
| **N-ter-based feature** |  |  |  |  |  |  |
| N-ter frequency-120d | 0.768±0.060 | 0.738±0.038 | 0.777±0.086 | 0.488±0.070 | 0.583±0.046 | 0.455±0.063 |
| N-ter frequency-200d | 0.905±0.021 | 0.799±0.057 | 0.934±0.028 | 0.772±0.075 | 0.782±0.042 | 0.724±0.054 |
| **pre-trained protein language-based feature** |  |  |  |  |  |  |
| ESM-2 | **0.939±0.012** | **0.845±0.061** | 0.964±0.018 | 0.870±0.058 | **0.855±0.027** | **0.818±0.033** |

Note: ACC: accuracy; SN: sensitivity; SP: specificity; PR: precision; F1: F1-score; MCC: Matthews correlation coefficient. The values were expressed as mean ± standard deviation. For each metric, the best performance value across different encoding methods is highlighted in bold.

**Supplementary Table S4.** Performance of various classifiers in this study evaluated by the independent test.

| Encoding | ACC | SN | SP | PR | F1 | MCC |
| --- | --- | --- | --- | --- | --- | --- |
| **PSSM-based feature** |  |  |  |  |  |  |
| AAC-PSSM | 0.906 | 0.929 | 0.903 | 0.591 | 0.722 | 0.694 |
| AADP-PSSM | 0.878 | 0.929 | 0.870 | 0.520 | 0.667 | 0.637 |
| AATP | 0.883 | 0.929 | 0.876 | 0.531 | 0.675 | 0.646 |
| AB-PSSM | 0.869 | 0.929 | 0.859 | 0.500 | 0.650 | 0.620 |
| DP-PSSM | 0.892 | 0.893 | 0.892 | 0.556 | 0.685 | 0.650 |
| EDP | 0.883 | 0.893 | 0.881 | 0.532 | 0.667 | 0.631 |
| EEDP | 0.925 | 0.893 | 0.930 | 0.658 | 0.758 | 0.726 |
| DPC-PSSM | 0.883 | 0.893 | 0.881 | 0.532 | 0.667 | 0.631 |
| k-separated-bigrams-PSSM | 0.911 | 0.929 | 0.908 | 0.605 | 0.732 | 0.704 |
| MEDP | 0.906 | 0.929 | 0.903 | 0.591 | 0.722 | 0.694 |
| Pse-PSSM | 0.906 | 0.964 | 0.897 | 0.587 | 0.730 | 0.707 |
| PSSM-AC | 0.850 | 0.893 | 0.843 | 0.463 | 0.610 | 0.572 |
| RPSSM | 0.915 | 0.964 | 0.908 | 0.614 | 0.750 | 0.728 |
| PSSM-composition | 0.878 | 0.857 | 0.881 | 0.522 | 0.649 | 0.606 |
| RPM-PSSM | 0.873 | 0.929 | 0.865 | 0.510 | 0.658 | 0.628 |
| S-FPSSM | 0.854 | 0.821 | 0.859 | 0.469 | 0.597 | 0.547 |
| smoothed-PSSM | 0.864 | 0.821 | 0.870 | 0.489 | 0.613 | 0.564 |
| TPC | 0.761 | 0.750 | 0.762 | 0.323 | 0.452 | 0.376 |
| tri-gram-PSSM | 0.798 | 0.571 | 0.832 | 0.340 | 0.427 | 0.329 |
| PSSM-CC | 0.901 | 0.821 | 0.914 | 0.590 | 0.687 | 0.642 |
| **Amino acid composition-based feature** |  |  |  |  |  |  |
| AAC | 0.817 | 0.679 | 0.838 | 0.388 | 0.494 | 0.415 |
| DPC | 0.770 | 0.714 | 0.778 | 0.328 | 0.449 | 0.368 |
| **N-ter-based feature** |  |  |  |  |  |  |
| N-ter frequency-120d | 0.831 | 0.964 | 0.811 | 0.435 | 0.600 | 0.577 |
| N-ter frequency-200d | 0.920 | 0.964 | 0.914 | 0.628 | 0.761 | 0.739 |
| **pre-trained protein language-based feature** |  |  |  |  |  |  |
| ESM-2 | **0.944** | **0.966** | **0.941** | **0.718** | **0.824** | **0.803** |

Note: ACC: accuracy; SN: sensitivity; SP: specificity; PR: precision; F1: F1-score; MCC: Matthews correlation coefficient. For each metric, the best performance value across different encoding methods is highlighted in bold.

**Supplementary Figure S1.**
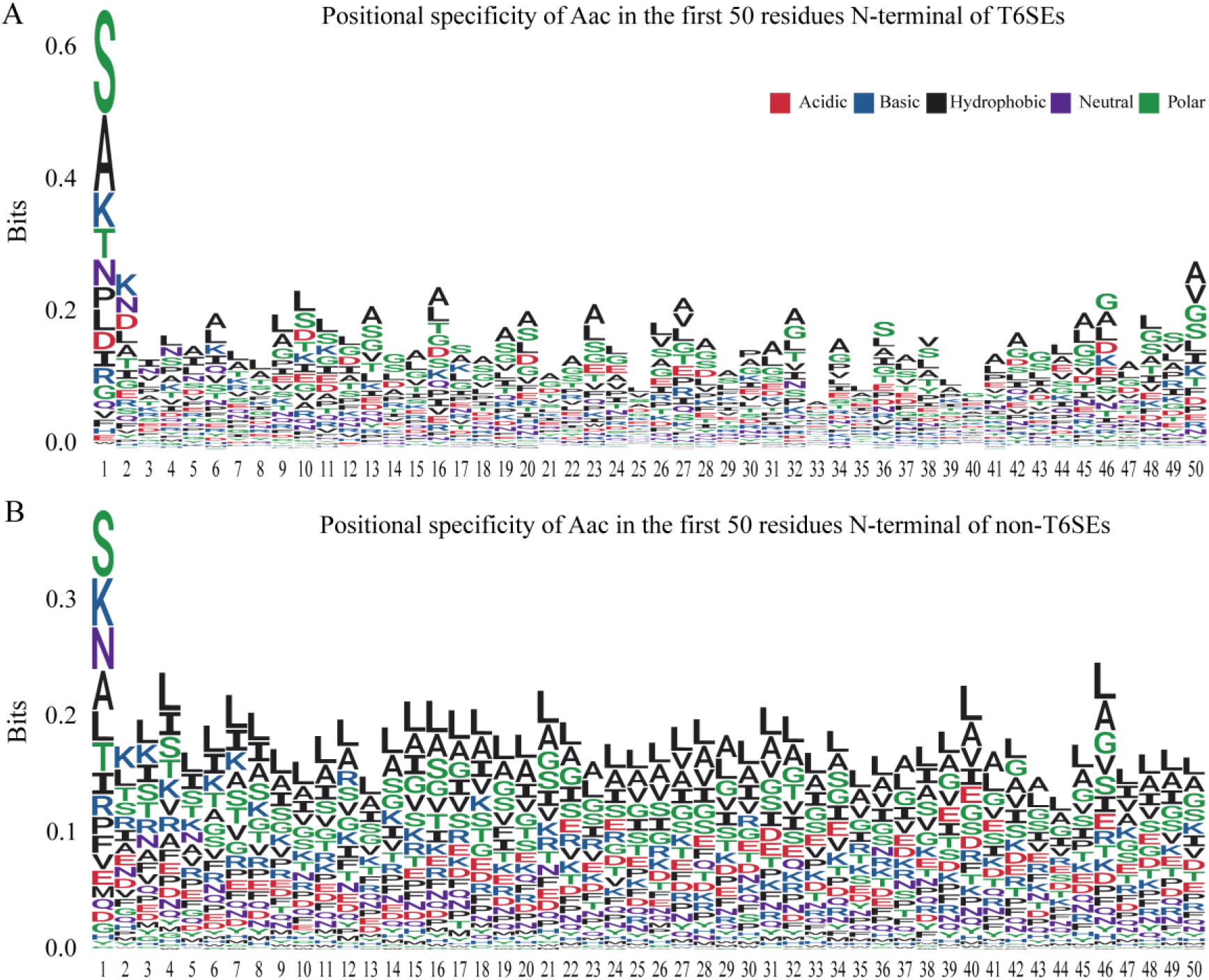
Positional specificity of amino acid composition in the N-terminal regions of T6SEs and non-T6SEs. (A) Positional specificity of amino acid composition in the first 50 residues of the N-terminal region of T6SEs. (B) Positional specificity of amino acid composition in the first 50 residues of the N-terminal region of non-T6SEs. The x-axis represents the position in the N-terminal sequence, while the y-axis indicates the information content at each position, in bits. The first position, typically occupied by methionine (M) in most proteins, is excluded. The stacked height at each position reflects the level of conservation, with the size of the letters representing the relative frequency of each amino acid.

**Supplementary Figure S2.**
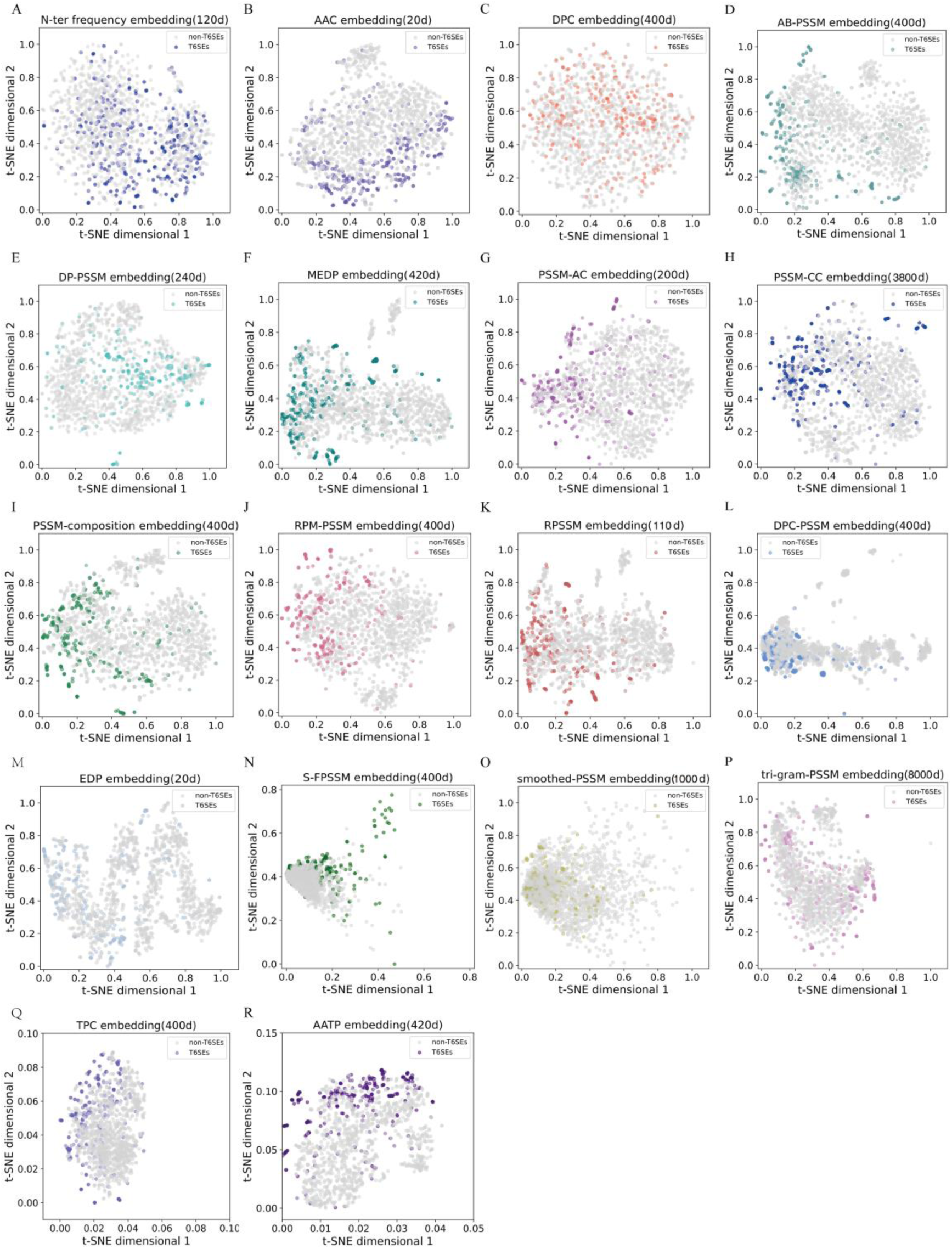
t-SNE visualization of embedding vectors generated by different feature encoding strategies. (A-R) shows the distribution of positive (colored) and negative (gray) samples after PCA dimensionality reduction, with each point representing a sample. The color coding indicates the distribution of positive samples under different feature encoding strategies.

**Supplementary Figure S3.**
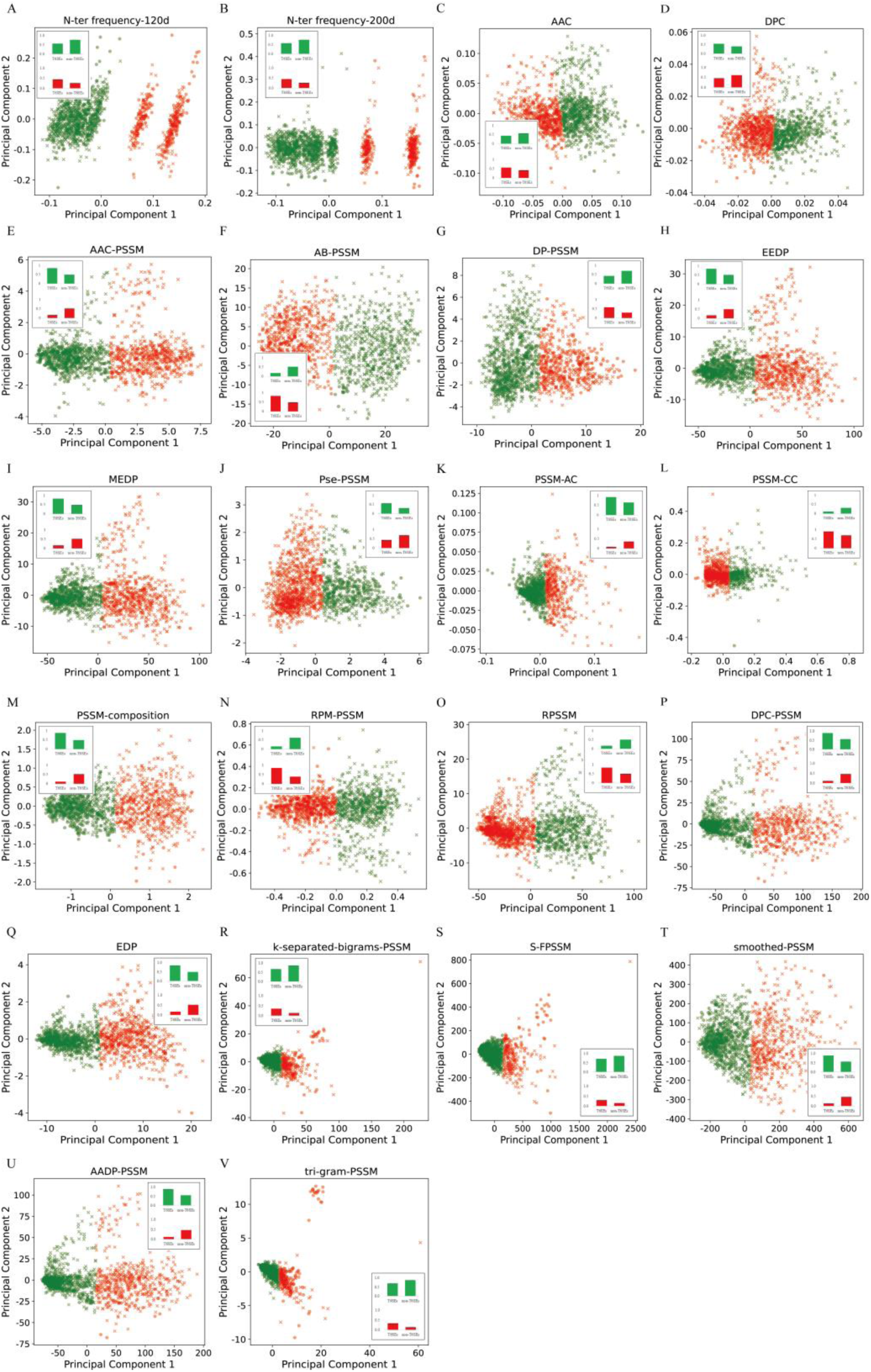
PCA representation and K-means clustering of T6SEs and non-T6SEs samples using different feature encoding strategies. For each feature encoding, samples are reduced to two dimensions using principal component analysis (PCA). K-means clustering is then applied to group the samples into two clusters, represented by green and red, with circles and crosses denoting T6SEs and non-T6SEs, respectively. The bar plots show the relative proportions of T6SEs (left) and non-T6SEs (right) within each cluster.

